# Topically applied thyroid hormones stimulate hair growth in organ-cultured human scalp skin

**DOI:** 10.1101/2024.06.11.598522

**Authors:** Jennifer Gherardini, Aysun Akhundlu, Matthew Gompels, Andrew Verbinnen, Sergi Velasco, Ulrich Knie, Ramtin Kassir, Jeremy Cheret, Ralf Paus

**Affiliations:** CUTANEON – Skin & Hair Innovations, Hamburg & Berlin, Germany; Dr. Phillip Frost Department of Dermatology & Cutaneous Surgery, University of Miami Miller School of Medicine, Miami, FL, USA; HairDAO, Palm Beach, FL, USA; Kassir Plastic Surgery, New York, NY, USA

## Abstract

We have previously shown that the thyroid hormones triiodothyronine (T3) and thyroxine (T4) prolong anagen, mitigate stem cell apoptosis, and stimulate mitochondrial functions in microdissected human scalp HFs ex vivo. To circumvent the systemic adverse effects of T3/T4, we have asked in the current pilot study whether topically applied T3/T4 retains hair growth-promoting properties. To prove this, we have topically treated healthy full-thickness human scalp skin with T3 (1, 10nM) and T4 (1, 10μM) for six days in serum-free organ culture, using an HF-targeting vehicle that contains only FDA-approved ingredients. This showed that, at distinct doses, topical T3 and T4 significantly increased the percentage of HFs in anagen, decreased the percentage of proliferative (Ki-67+) cells in the hair matrix, did not promote melanogenesis (as measured by quantitative Warthin-Starry histochemistry), and significantly increased keratin 15 expression in the bulge. Finally, T3 and T4, at low concentrations, increase the expression of the hair growth promoters IGF-1 and FGF-7. The lower concentration of T3 and both of T4 also significantly increases the number of CD31+ endothelial cells, suggesting a pro-angiogenic effect, which is also important for hair growth promotion. These preliminary results strongly suggest that topically applied thyroid hormones promote hair growth in intact human scalp on multiple levels ex vivo. This invites the intermittent pulse application of topical T3 and T4 as a novel therapeutic intervention for managing hair loss disorders associated with telogen effluvium, such as androgenetic alopecia.

## INTRODUCTION

Clinically, thyroid hormones (THs), namely triiodothyronine (T3) and L-thyroxine (T4), have long been appreciated as endocrine mediators whose serum levels potently affect human hair growth and hair shaft quality (van Beek et al. 2008; Gharaei Nejad et al. 2022; Hussein et al. 2023). Yet, THs remain to be systematically explored as a candidate for hair growth-modulatory therapeutics in dermatology (Hussein et al. 2023; Paus et al. 2020). These nuclear hormone receptor agonists primarily, but by no means only, signal via forming a complex with the nuclear TH receptors c-erb Aα and Aβ (TRα/β), which then preferentially binds to the promoter region of genes with a TH response element, thus regulating the transcription of a vast number of target genes, including in human skin and hair follicles (HFs) (Brtko 2021; Paus et al. 2020; Penna et al. 2023). From a hair growth perspective, THs deserve scrutiny for the following reasons:

THs alter hair growth *in vivo* in mice, rats, sheep, and humans (Hale and Ebling 1975; Hussein et al. 2023; Safer et al. 2001). Rapidly growing (anagen) human scalp HFs prominently express TRβ, can enzymatically deiodinate T4 into T3, and directly respond to TH stimulation *ex vivo* (van Beek et al. 2008; Kaplan et al. 1988; Safer et al. 2009). Human anagen scalp HFs are unusually sensitive to even minor variations in the serum level of the two main THs, T3, and T4 - independent of whether these are disease-related or caused by medication with T4, one of the most frequently prescribed drugs in clinical medicine (van Beek et al. 2008; Hussein et al. 2023). In patients with hypothyroidism, this can result in substantially increased hair shaft shedding (telogen effluvium) and brittle, dull, and dry hair shafts (Freinkel and Freinkel 1972; Schell et al. 1991). Paradoxically, chronically hyperthyroid patients can show very similar hair pathology, although - besides telogen effluvium - a thinner hair shaft diameter, reduced tensile strength, and greasy hair tend to predominate here (Fistarol 2002; Messenger 2000; Stüttgen and Schaefer 1974).

Yet, both T3 and T4 stimulate at different concentrations the proliferation of hair matrix keratinocytes, alter the intrafollicular production of selected keratins, and prolong the duration of the growth stage of the hair cycle, anagen(van Beek et al. 2008; Paus 2010; Schell et al. 1991; Vidali et al. 2014). The latter is also seen when human scalp HFs are treated *ex vivo* with eprotirome, a synthetic TR agonist that signals mainly via TR-β (Oláh et al. 2016; Shiohara et al. 2012). Furthermore, THs stimulate mitochondrial activity and likely biogenesis in the hair matrix (Vidali et al. 2014). This renders THs attractive candidate agents that may effectively counter telogen effluvium, which results from the premature induction of catagen in anagen HFs (Chien Yin et al. 2021; Harries et al. 2016; Rebora 2019). An inhibitory effect of THs on telogen effluvium would also be of significant clinical interest in the management of male pattern androgenetic alopecia and female pattern hair loss, both of which can be associated with prominent telogen effluvium (Chien Yin et al. 2021; Harries et al. 2016; Rebora 2019).

The well-known, considerable toxicity of increased serum levels of TH (Lee and Pearce 2023) renders it both prudent and responsible to apply these potent peptide hormones only topically to a limited (scalp) skin surface area to sharply reduce the risk of systemic adverse effects and the induction of feedback counter-regulatory mechanisms in the central hypothalamus-pituitary-thyroid (HPT) axis that regulates systemic TH levels (Braverman and Cooper 2012; Fekete and Lechan 2014). However, it remains entirely unknown whether topically applied TH can penetrate the relatively thick epidermal barrier of human scalp skin *and* reach sufficient bioactive hormone levels to impact in a clinically desirable manner on anagen hair bulbs located deep in human scalp skin, i.e., in the upper subcutis (Oh et al. 2016; Poblet et al. 2018). The current *ex vivo* pilot study aimed to probe whether this is possible in principle.

Specifically, we asked whether topical application of T3 or T4 or combined treatment with both can prolong anagen in human scalp HFs that grow within healthy, organ-cultured adult scalp skin, using our established scalp skin organ culture assay(Alam et al. 2020; Bertolini et al. 2023; Gherardini et al. 2019; Vidali et al. 2016). Therapeutic anagen prolongation, which is indistinguishable from catagen inhibition, is the crucial prerequisite for countering telogen effluvium since the latter is caused by premature anagen termination and catagen induction (Chien Yin et al. 2021; Paus and Cotsarelis 1999; Rebora 2019).

After preparatory exploration of different vehicle compositions, we opted for two distinct ethanol- and propylene glycol-based vehicles that contain only FDA-licensed ingredients, promised to maximize THs penetration under skin organ culture conditions, and differed in their relative ethanol and propylene glycol content (details, see below). Two TH doses that we had previously shown to stimulate the growth of microdissected, isolated scalp HFs (van Beek et al. 2008) and to promote wound healing of human skin ex vivo (Post et al. 2021; Zhang et al. 2019) were applied topically every other day to human scalp skin, organ-cultured at the air-liquid interphase in supplemented, serum-free medium (Bertolini et al. 2023; Gherardini et al. 2019), i.e. in the absence of THs in the medium and of any confounding systemic or neural inputs.

Besides studying the impact of TH on the percentage of anagen HFs by quantitative hair cycle histomorphometry, we also assessed hair shaft production *ex vivo* as well as melanin content, hair matrix keratinocyte proliferation/apoptosis in anagen hair bulbs, and intrafollicular protein expression of the critical hair growth-regulatory growth factors, IGF-1, TGF-β2, and FGF-7/KGF by quantitative (immuno-)histomorphometry as described previously (Alam et al. 2020; Bertolini et al. 2023; Gherardini et al. 2019).

Since we had previously noted potentially undesirable effects of ‘systemically’ applied THs on keratin 15+ human HF epithelial stem cells in isolated, microdissected, and therefore experimentally traumatized scalp HFs *ex vivo* (Tiede et al. 2010), we investigated intrafollicular protein expression *in situ* of the key stem cell-associated keratin, K15, after topical TH treatment of intact, organ-cultured human scalp skin. Finally, we also assessed whether dermal CD31+ endothelial cells responded to topical THs stimulation *ex vivo*, given that we had previously found ‘systemically’ applied T4 to enhance CD31 expression and possibly angiogenesis in experimentally wounded, organ-cultured human skin (Post et al. 2021; Zhang et al. 2019). This would be expected to exert indirect hair growth-promoting effects (Cheng et al. 2017; Mecklenburg et al. 2000; Yano et al. 2001) and could help prevent the blood vessel regression from the dermal papilla of HFs from the balding skin area of androgenetic alopecia patients (Deng et al. 2022).

The current pilot study provides the first proof of principle that topically applied TH can promote hair growth at different levels of hair physiology in intact human scalp skin *ex vivo*.

## MATERIALS & METHODS

### Human skin sourcing

Frontotemporal and occipital human scalp skin specimens from healthy female and male patients undergoing facelift surgery were collected for this study. These human discarded tissues are considered non-human subject research and exempted under 45 CFR46.101.2 by the Institutional Review Board of the University of Miami Miller School of Medicine (Miami, FL). Samples used for this study were taken from 5 different and independent donors (age range 48-56 y-o, mean age 51 years), were obtained one day after surgery, and 4 mm skin fragments were performed.

### Human scalp skin organ culture

4 mm skin fragments were performed and placed at an air-liquid interface in a cell strainer (with the subcutis and dermis located in the medium to avoid that epidermis falling into the culture medium during the culture) within serum-free supplemented William’s E media and incubated at 37°C in a humidified atmosphere of 5% CO_2_. After one day of culture for equilibration, 4 mm skin fragments were treated topically with 2 μl of viscous formulation A or B prepared in our laboratory (hydroxypropyl cellulose was added to increase viscosity and prevent test compound spill-over from the skin surface into the medium) either containing William’s E medium for the vehicle, or T3 (1 or 10nM) or T4 (1 or 10μM), for five days. Medium change and topical administration of test compounds (any leftover test compound was removed before applying fresh T3/T4) were performed every other day (see Table 1). At Day 0, 1, 6, and during every medium change, the surface of each 4 mm skin fragment was imaged to perform hair shaft production analysis. At the end of the six days of culture, samples were embedded in OCT and snap-frozen in liquid nitrogen before being stored at −80°C until further analyses.

**Table 1.** Experimental design.

| Day 0 | Day 1 | Day 2 | Day 3 | Day 4 | Day 5 | Day 6 |
| --- | --- | --- | --- | --- | --- | --- |
| Perform 4 mm skin fragments<br>5ml of supplemented medium/well | Take picture for hair shaft production analysis<br>Change medium/Add treatment :<br>• Vehicle<br>• T3 1 nM topical A/B<br>• T3 10 nM topical A/B<br>• T4 1 uM topical A/B<br>• T4 10 uM topical A/B<br>• T3 10nM + T4 10uM (Combo) A/B | Rest | Take picture for hair shaft production analysis<br>Change medium/Add treatment :<br>• Vehicle<br>• T3 1 nM topical A/B<br>• T3 10 nM topical A/B<br>• T4 1 uM topical A/B<br>• T4 10 uM topical A/B<br>• T3 10nM + T4 10uM (Combo) A/B | Rest | Take picture for hair shaft production analysis<br>Change medium/Add treatment :<br>• Vehicle<br>• T3 1 nM topical A/B<br>• T3 10 nM topical A/B<br>• T4 1 uM topical A/B<br>• T4 10 uM topical A/B<br>• T3 10nM + T4 10uM (Combo) A/B | Take picture for hair shaft production analysis<br>Freeze punches in OCT |

### Vehicles and TH doses tested

#### Vehicle A

Vehicle A consisted of 30% Ethanol (v/v), 5% hydroxypropyl cellulose (w/v), 50% propylene glycol (v/v), and purified water to 1 ml.

#### Vehicle B

Vehicle B consisted of 60% Ethanol (v/v), 5% hydroxypropyl cellulose (w/v), 20% propylene glycol (v/v), and purified water to 1 ml.

Since the current study only aimed to investigate whether or not it is possible to manipulate human hair growth-associated key read-out parameters *ex vivo* by topically applied THs, vehicle design was primarily guided by maximizing the chance of reaching a bioactive TH concentration in anagen hair bulbs within intact, organ-cultured human scalp skin, using as few FDA-approved vehicle components as possible in the presence of water to reduce the risk of deleterious skin desiccation by excessively high alcohol content. A further requirement was to design formulations that could be prepared easily at the test-site lab without sophisticated technologies. Plain solvent mixtures were the first choice for vehicles. Ethanol, isopropanol, propylene glycol, and water were suitable solvents, but ethanol was preferred to isopropanol due to its better smell and organic properties. Propylene glycol is extensively used in numerous topical formulations (WILLIAMS 2007) as a solvent and co-solvent. Propylene glycol has a penetration-enhancing effect for many active ingredients, which can be attributed to its solvent properties and the interaction with skin lipids and proteins (Carrer et al. 2020; Yu and Goh 2024). For curcumin nanocrystals, for example, propylene glycol allows for hair follicle targeting and intense passive dermal penetration simultaneously (Pelikh and Keck 2020). Hydroxypropyl cellulose was added to increase the viscosity of the vehicle. Solvent mixtures of ethanol/propylene glycol/water are used to formulate a well-known topically applied hair growth promoter, Minoxidil. For topically applied products, one must remember the so-called metamorphosis (Surber and Knie 2018). That means the sum of vehicle ingredients remaining on the skin will change dramatically after application onto the skin compared to the original formulation due to the evaporation of the volatile ingredients or absorption of ingredients into the skin. In the case of the proposed ethanol/propylene glycol/water mixture, a great deal of ethanol and water will evaporate or be absorbed into the skin, leaving behind a solution of T3 and/or T4 in a residue consisting mainly of propylene glycol.

The concentration of THs in this residual vehicle is critical because it dramatically impacts the flux of THs through the epidermal barrier. The flux is directly proportional to the thermodynamic activity of the active ingredients in the residual vehicle, which is related to the degree of saturation in the vehicle. Concerning the proposed formulations, the metamorphosis phenomenon would nearly double the concentration of the THs in the residual vehicle of formulation A and almost quintuple in all the residual of formulation B. Consequently, although the concentrations of THs are the same in the primary formulations, the flux produced with formulation B should be higher.

While formal pharmaceutical development, as needed for human clinical studies, was not executed, it deserves notice that T3 and T4 are stable in unbuffered aqueous systems for oral use (Bellorini et al. 2013; Parikh and Hite 2016). Therefore, topical test solutions freshly prepared before application on scalp skin organ can be justified without long-term stability studies. The concentrations of THs (1 and 10nM for T3; 1 and 10µM for T4) have been chosen based on our previous studies on human scalp HF organ cultures (van Beek et al. 2008; Hardman et al. 2015a; Post et al. 2021; Tiede et al. 2010; Vidali et al. 2016; Vidali et al. 2014; Zhang et al. 2019) where biological effects using “systemic” application of these THs to the culture medium were observed. For topical application, we opted to test 10 times higher concentrations than the most efficient concentrations observed in our previous “systemic” application ex vivo work to maximize the chance that effective TH concentrations are reached within the anagen hair bulb in intact scalp skin.

### Melanin histochemistry

Warthin-Starry histochemistry was carried out to assess the melanin content of the HFs according to the previously established protocol (Joly-Tonetti et al. 2016; Lai and Healy 2016; Samra et al. 2023; Sevilla et al. 2023). Briefly, slides were fixed in 4% paraformaldehyde and then washed with PBS and distilled water. The cryosections were next stained with 0.5% silver nitrate and incubated at 50 °C for 6 minutes. A reducing agent solution of 4% acidulated gelatin, 2% acidulated silver nitrate, and 0.1% hydroquinone solution was added to each section and incubated at 50 °C for 5 minutes. The sections were washed in PBS, counterstained with hematoxylin, and dehydrated in 100% 2-propanol before embedding with VectaMount Express Mounting Medium (Vector Laboratories).

### Hair cycle staging and hair cycle score

Masson–Fontana histochemistry and Ki-67/Caspase-3 immunostainings were used for accurately determining hair cycle staging *ex vivo*, using the well-defined classification criteria described in detail (Kloepper et al. 2010; Oh et al. 2016).

### Immunofluorescence staining

OCT-embedded HFs were cryosectioned (7 μm thickness) with a Crysostar NX50 (Thermo Scientific, Epredia) cryostat. For immunofluorescence staining, all samples/sections were fixed and pre-incubated for 30 minutes at RT, followed by incubation with a primary antibody (previously used and characterized; see references in table 2) overnight at 4°C. After three washes for five minutes each, sections were incubated with a fluorescent-tagged secondary antibody for 45 minutes at room temperature and embedded in DAPI/Fluoromount (Electron Microscopy Sciences). All details about fixatives, blocking reagents, and primary and secondary antibodies used in this study are listed in **Table 2**. Negative control for the primary antibody was used by omitting the primary antibody.

**Table 2.**
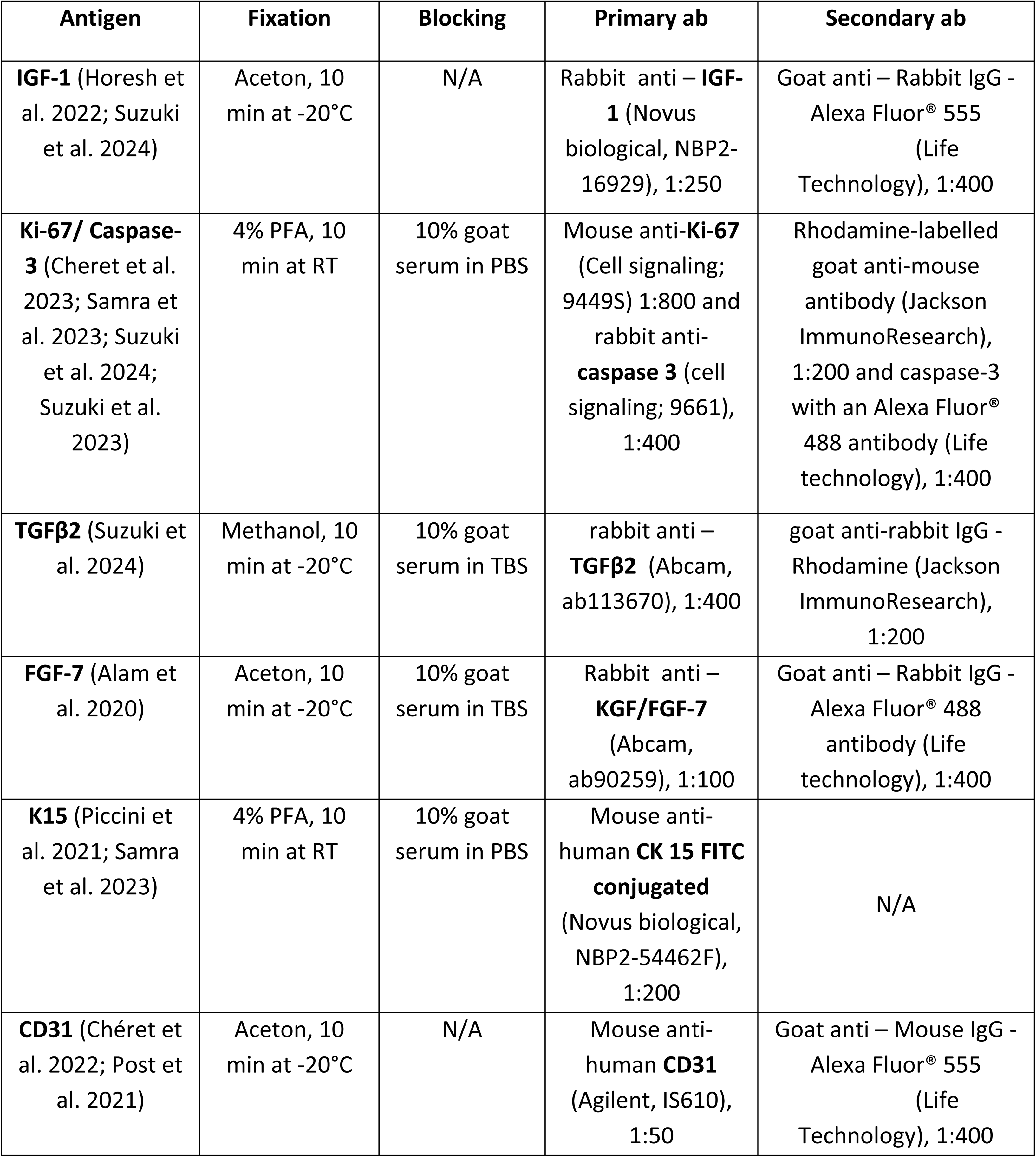
Immunofluorescence microscopy: Details.

### Quantitative immunohistomorphometry and microscopy

Images were taken using a BZ-X800 All-in-one Fluorescence Microscope (Keyence Corporation) and Image Analysis software Bz-800 Analyzer (Keyence Corporation) at 200x magnification, with a constant exposure time used throughout imaging to allow for later normalization. Images were then analyzed via NIH ImageJ software (National Institute of Health), using defined reference areas (dotted area shown in the vehicle images). For staining intensity, the mean intensity was measured. The ImageJ cell counter function was used for single and double-positive cell counts.

### Statistical analysis

All data are presented as fold change of the mean ± SEM. We employed the Student’s t-test or the Mann-Whitney test for statistical analysis, depending on whether the data adhered to a Gaussian or non-Gaussian distribution, respectively (as determined by the d’Agostino and Pearson omnibus normality test). This analysis used Graph Pad Prism 9 software (GraphPad Software, San Diego, CA, USA). Statistical significance was set at p<0.05.

## RESULTS

### Topically applied T3 1nM significantly increases the percentage of anagen VI hair follicles in intact human scalp skin

It is well-known that hair shaft production in organ-cultured human scalp skin is much slower, more heterogeneous, and more unpredictable than in microdissected, amputated scalp HFs *ex vivo* (Langan et al. 2015; Lu et al. 2007). Therefore, we did not expect noticeable effects of topical THs application on hair shaft production during the relatively short time of organ culture (6 days) and examined the percentage of HFs that visibly generated a longer hair shaft over the course of organ culture, which demonstrates that these HFs were in anagen, only in selected skin fragments. Though our minimal sample size does not permit reliable conclusions, the obtained macroscopic data already raised the possibility that topical T3 (namely 1nM) in vehicle A may keep more HFs in anagen compared to vehicle-treated control samples (p=0.067) and all other test groups **(Suppl Fig. S1A-D)**. This cursory initial macroscopic observation was systematically followed up histologically.

Quantitative hair cycle histomorphometry (Kloepper et al. 2010; Oh et al. 2016) showed that, when vehicle A was used, topical T3 1nM significantly increased the percentage of anagen HFs in organ-cultured human scalp skin of three independently tested donors **(Fig. 1A, C)**, corroborating the initial hair shaft production observations **(Suppl Fig. S1A-B)**. Higher doses of T3 (10nM) and T4 (10µM) also showed a higher percentage of HFs in anagen, even though significance was not reached. Unexpectedly, the T3 10nM +T4 10µM combination had the opposite effect in vehicle A and may even promote premature catagen induction **(Suppl Fig. S1E)**. However, in vehicle B, this TH combination was not hair growth-inhibitory **(Suppl Fig. S1F)**. The higher doses of T3 and T4 individually tended to increase the % of anagen HFs **(Fig. 1D-F)**, but the differences were again not statistically significant.

**Figure 1:**
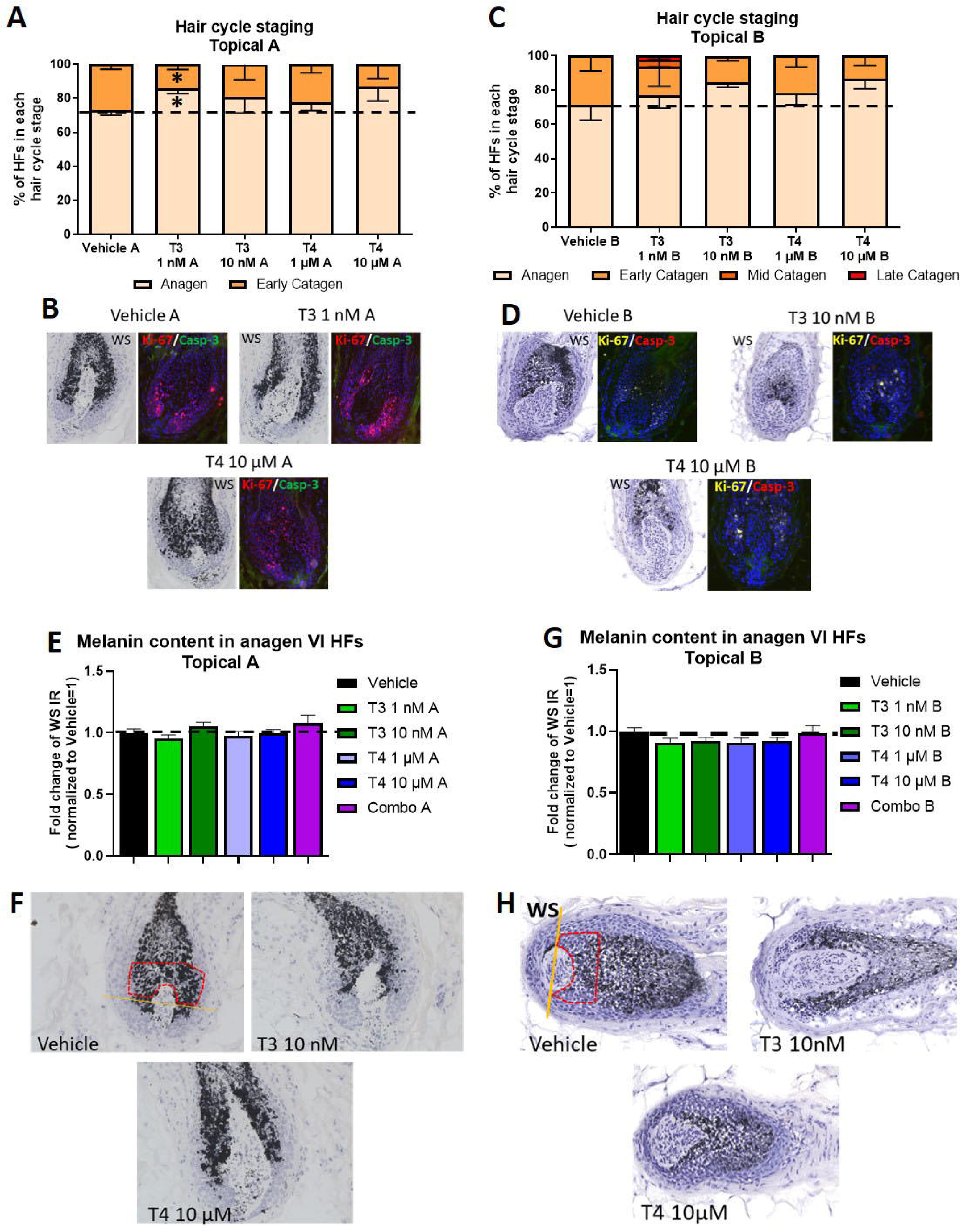
Topical T3 1nM significantly prolongs anagen while topical thyroid hormones do not affect melanin production in human scalp skin *ex vivo*. **(A-C)** Hair cycle staging was performed using Ki-67 and Masson-Fontana histochemistry as described (Kloepper et al. 2010; Oh et al. 2016). Mean +/- SEM; n=20-35 HFs from 3 donors treated with T3 (1 or 10nM), T4 (1 or 10 µM), or vehicle; Mann–Whitney test, *p<0.05. **(B, D)** Representative fluorescence images of Ki-67 and bright-field microscopic images of Masson-Fontana histochemistry, used for accurate hair cycle staging. **(E, G)** Quantitative immunohistomorphometry (qIHM) of melanin production. Mean +/- SEM; n=15-33 HFs from 3 donors treated with T3 (1 or 10nM), T4 (1 or 10 µM), or vehicle; Mann–Whitney test, not significant. **(F, H)** Representative bright field microscopic images of Masson-Fontana histochemistry. Red dotted area: reference areas in the hair bulb of non-consecutive sections. Orange line: Auber’s line. Magnification: 200x.

When HFs were compared with each other by Ki-67/caspase-3 quantitative immunohistomorphometry (qIHM), the above hair cycle effects of the TH did not translate into a significant stimulation of hair matrix keratinocyte proliferation by the end of scalp skin organ culture **(Suppl Fig. S1G-J)**. This is not surprising, as test agents typically exert significant effects on hair matrix proliferation during the first few days of organ culture (Kloepper et al. 2010), which could not be assessed separately due to the very limited availability of scalp skin samples. Furthermore, our topical T3 data with vehicle A and B align with the hair growth-promoting effect we had previously seen with “systemic” application (van Beek et al. 2008). In line with our macroscopic hair shaft observations **(Suppl Fig. S1A-D)**, the combination of both topically applied THs even significantly inhibited hair matrix proliferation *ex vivo* in vehicle A compared to the relevant vehicle control **(Suppl Fig. S1E)**.

Of note, no apoptotic hair matrix keratinocytes were seen in any of the test or control groups using either vehicle A or B (data not shown). Together with the many Ki-67+ cells seen in the hair matrix in all test and control skin fragments, this attests to the continued viability of scalp skin for the entire duration of organ culture.

### Topical THs do not significantly promote intrafollicular melanin production in scalp skin *ex vivo*

Next, we investigated the impact of topical THs on HF pigmentation, as assessed by particularly sensitive Warthin-Starry melanin histochemistry (Joly-Tonetti et al. 2016; Sevilla et al. 2023) (Joly-Tonetti et al. 2016; Sevilla et al. 2023a). Since even slight abnormalities in the distribution pattern and size of melanin granules within the anagen hair bulb are a very sensitive indicator of HF damage (Bodó et al. 2007; Cheret et al. 2023; Samra et al. 2023; Suzuki et al. 2023), it is important to note that no melanin clumping or ectopic melanin granules were observed in either of the test groups independent of the vehicle employed and whether the THs were applied alone or in combination **(Fig. 1F, H)**. Together with the absent induction of hair matrix apoptosis (see above), this indicates that topical THs are well-tolerated by the anagen HF and its highly damage-sensitive pigmentary unit (Bodó et al. 2007; O’Sullivan et al. 2021).

We did not observe a significant increase in HF melanin production by topically applied THs during scalp skin organ culture **(Fig. 1E-H)**. This contrasts with the substantial stimulation in intrafollicular melanogenesis by “systemic” TH application we had previously seen in microdissected, organ-cultured HFs ex vivo, namely of T4 (van Beek et al. 2008; Paus et al. 2020), possibly due to a much lower % of topically TH applied in an aqueous vehicle that reaches the HF pigmentary unit compared to adding TH directly to the organ culture medium.

### Topical THs significantly enhance intrafollicular FGF-7 and IGF-1 production

Next, we interrogated the impact of topical THs on the intrafollicular production of three key hair growth-modulatory growth factors in human scalp HFs: the anagen-promoting growth factors, FGF-7 and IGF-1 (Alam et al. 2020; Booth and Potten 2000; Chéret et al. 2018; Philpott et al. 1994; Rudman et al. 1997), and the most critical physiological catagen-promoting growth factor, TGF-β2 (Foitzik et al. 2006; Hibino and Nishiyama 2004; Soma et al. 2002).

qIHM showed that topical application of both THs to intact human scalp skin *ex vivo* stimulated to some extent, and in combination significantly, IGF-1 protein synthesis in the outer root sheath (ORS) - in vehicle B more so than in vehicle A **(Fig. 2A-D)**. In vehicle A, both THs and the combination also significantly promoted FGF-7 protein production in the ORS **(Fig. 2E-F),** while that of TGF-β2 was unaffected **(Suppl Fig. S2A-B)**. Though less pronouncedly, the THs also increased FGF-7 production when administered in vehicle B **(Fig. 2G-H)**. In this vehicle, the lower tested dose of T4 also significantly upregulated TGF-β2 production **(Suppl Fig. S2C-D),** which does not recommend this preparation for hair growth promotion. Overall, these preliminary data raise the possibility that the anagen-prolonging effects of topical THs are mediated, at least in part, by up-regulating the production of FGF-7 and, much less prominently, of IGF-1 within the HF epithelium in intact human scalp skin.

**Figure 2:**
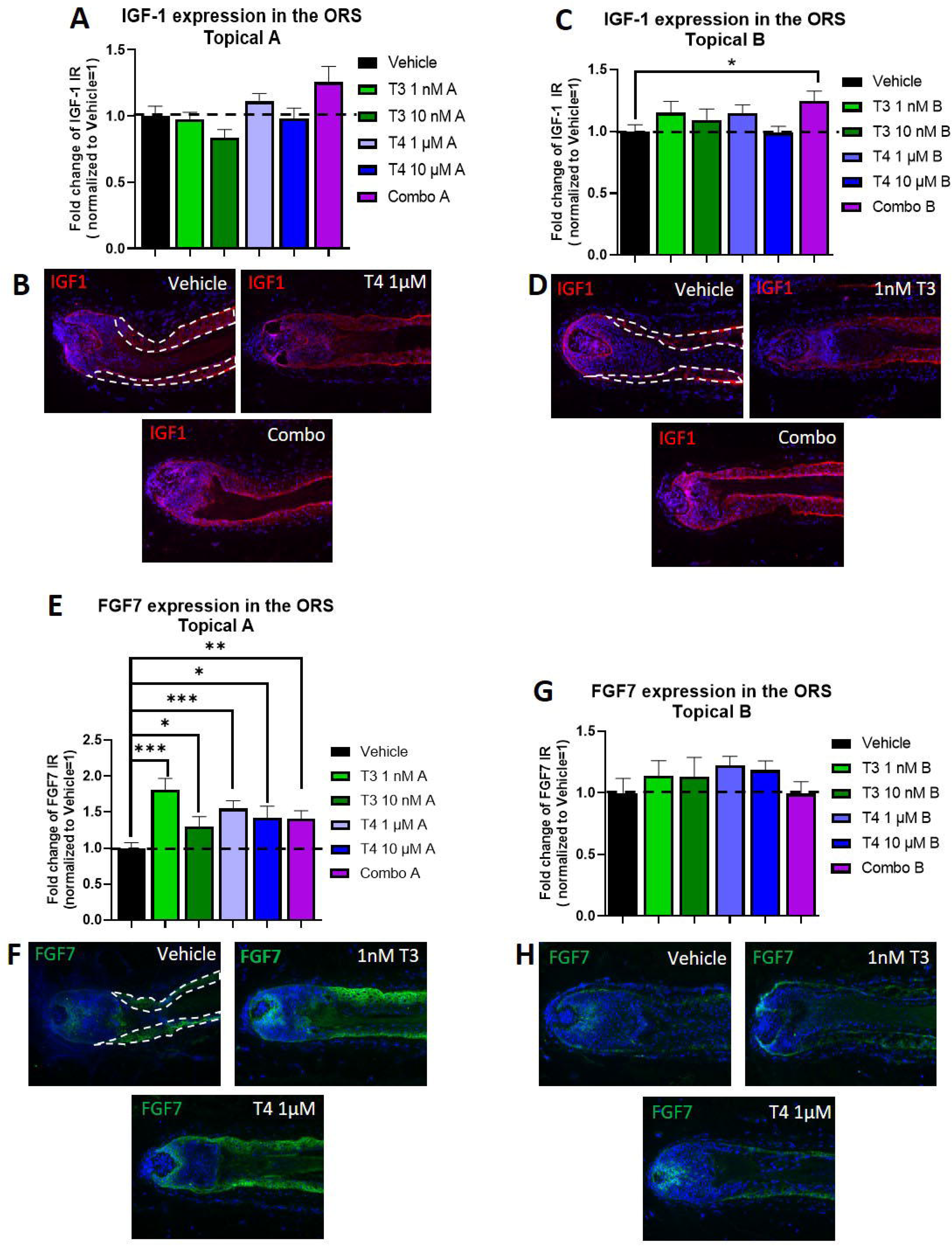
Topical thyroid hormones significantly increase protein expression of the anagen-prolonging growth factors, IGF-1 and FGF-7, in human scalp skin ex vivo. **(A, C)** qIHM of IGF-1 expression. Mean +/- SEM; n=18-35 HFs from 3 donors treated with T3 (1 or 10nM), T4 (1 or 10 µM), or vehicle; Mann–Whitney test, *p<0.05. **(B, D)** Representative images of IGF-1 immunofluorescence. **(E, G)** qIHM of FGF-7 expression. Mean +/- SEM; n=17-33 HFs from 3 donors treated with T3 (1 or 10nM), T4 (1 or 10 µM), or vehicle; Mann–Whitney test, *p<0.05, **p<0.01, ***p<0.001. **(F, H)** Representative images of FGF-7 immunofluorescence. White dotted area: reference areas used for qIHM in the proximal outer root sheath of non-consecutive sections. Magnification: 200x.

### Topical THs increase keratin 15 expression in the bulge

We also performed preliminary analyses to probe whether topical THs deplete the HF’s fundamental niche of K15+ epithelial stem cells in the bulge—a possibility suggested by our earlier HF organ culture study (Tiede et al. 2010). qIHM of organ-cultured scalp skin from 3 donors treated with topical TH over five days showed that this either does not reduce the number of K15+ cells in the bulge or even increases their number and/or the intensity of K15 protein expression in the bulge **(Fig. 3A-F).**

**Figure 3:**
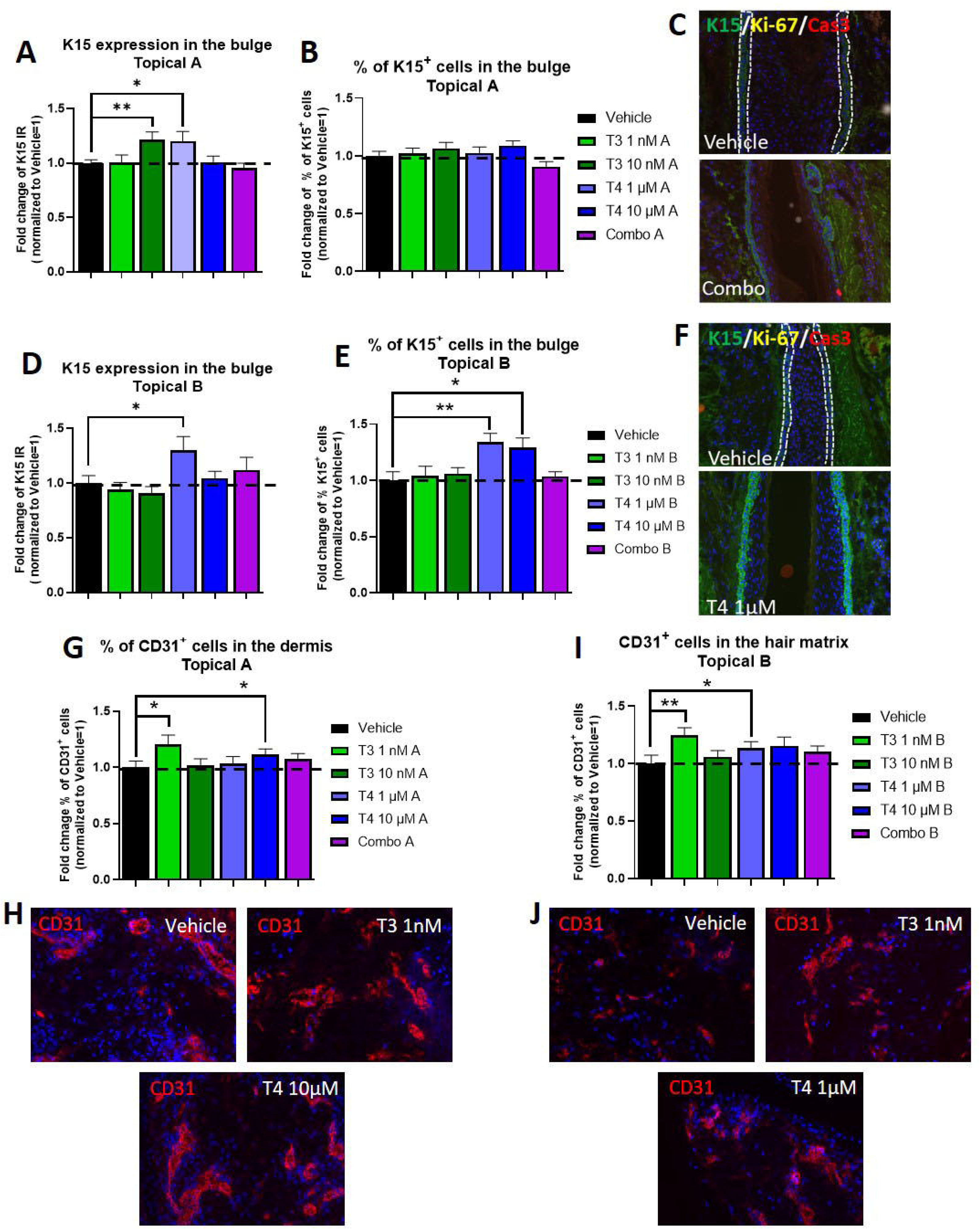
Topical thyroid hormones significantly increase keratin 15 expression in the bulge and the number of dermal endothelial cells in human scalp skin. **(A-B, D-E)** qIHM of K15 expression **(A, D)** and K15+ cell number **(B, E)**. Mean +/- SEM; n=21-34 HFs from 3 donors treated with T3 (1 or 10nM), T4 (1 or 10 µM), or vehicle; Mann–Whitney test, *p<0.05, **p<0.01. **(C, F)** Representative images of K15/Ki-67/Caspase-3 immunofluorescence. **(G, I)** qIHM of CD31+ cell number in the papillary dermis 200μm below the basal membrane. Mean +/- SEM; n=22-34 HFs from 3 donors treated with T3 (1 or 10nM), T4 (1 or 10 µM), or vehicle; Mann–Whitney test, *p<0.05, **p<0.01. **(H, J)** Representative images of CD31 immunofluorescence. White dotted area: reference areas used for qIHM in the bulge of non-consecutive sections. Magnification: 200x.

While topical T3 10nM and T4 1μM administered in vehicle A significantly up-regulated the latter, topical T4 in vehicle B even increased the percentage of K15+ cells detectable by IF microscopy in the bulge **(Fig. 3A, D-F),** rather than depleting the stem cell niche, at least during the six days of organ culture. This also further underscores that both selected topical vehicles deliver a substantial amount of bioactive THs to the scalp HF epithelium, including to the bulge, the HF’s stem cell niche.

### Topical THs also increase the number of dermal endothelial cells in human scalp skin

Since we had previously observed that TH may promote angiogenesis in experimentally wounded human skin ex vivo (Post et al. 2021; Zhang et al. 2019), we finally asked whether topically applied TH impact at all dermal endothelial cells in organ-cultured human scalp skin. Interestingly, the lower dose of topical T3 and T4 in vehicle A as well as T3 1nM and T4 10μM in vehicle B slightly but significantly increased the percentage of CD31+ endothelial cells in the dermis **(Fig. 3G-J)**. This also demonstrates that topically applied THs reach the dermis of human scalp skin *ex vivo* and are bioactive despite the use of water-containing vehicles.

## DISCUSSION

Our pilot study provides the first evidence in the literature that topically applied THs can prolong the duration of anagen even in intact human scalp skin *ex vivo*. Though our study can only be considered preliminary, since the low number of donors and scalp fragments that were available for study does not permit definitive conclusions, the current proof-of-principle is in line with our previous observation that TH promote the growth of microdissected, amputated HFs after administration to the culture medium, thus imitating a systemic mode of hormone application (van Beek et al. 2008). Our preliminary findings also correspond well to the hair growth stimulation that has previously been reported after topical T3 administration to mouse skin *in vivo* (Safer et al. 2003; Safer et al. 2001) - and this, even though very thin mouse epidermis is much more penetrable for numerous test agents than the thick, multi-layered and relatively penetration-resistant epidermal barrier of human scalp skin.

That both topical T3 and T4 prolonged anagen in human scalp skin *ex vivo* in both vehicles tested strongly suggests that these THs are likely to inhibit telogen effluvium in AGA patients. This would prolong the window during which miniaturized vellus HFs can be reconverted into terminal HFs, which is possible only during anagen (Gilhar et al. 2023; Gilhar et al. 2022; Pantelireis and Higgins 2018). The difference in effects we observed for the same TH at the same concentrations between the two topical vehicles tested here may result from the quantity of bioactive TH reaching defined HF compartments, corresponding to the metamorphosis phenomenon (see above), which would nearly double the concentration of THs in the residual vehicle of formulation A and almost quintuple it in the residuum of formulation B. Unexpectedly, our preliminary findings suggest that the combined application of T3 and T4, at least in the doses and vehicles tested here, promises no added benefit and may even exert undesirable hair growth-inhibitory effects. The underlying mechanisms remain speculative but might relate to the activation of alternative, non-classical TH signaling pathways by this combination and could reflect associated changes in the transcriptional regulation of different sets of target genes in human skin (Paus et al. 2020), depending on which dose of T3 or T4 is administered alone respectively in combination.

Functionally, our growth factor analyses suggest but do not yet prove, that the observed anagen-prolonging effects of topical T3 and T4 (alone) may be mediated, at least in part, through the stimulation of intrafollicular FGF-7 and, much less prominently, IGF-1 production - two recognized key hair growth-promoting growth factors in human scalp HFs (Alam et al. 2020; Booth and Potten 2000; Chéret et al. 2018; Philpott et al. 1994; Rudman et al. 1997). Such functional evidence could be obtained by repeating the reported experiments in the presence of FGF-7- or IGF-1-neutralizing antibodies (Alam et al. 2020; Chéret et al. 2018). Given the known stimulation of HGF and VEGF-A production by THs (Li et al. 2017; Zhang et al. 2010) and recognizing that these growth factors can also promote human hair growth (Li et al. 2012; Lindner et al. 2000; Nicu et al. 2021; Yano et al. 2001; Yoon et al. 2019), it also deserves to be explored whether increased HGF and/or VEGF-A production occurs after topical TH administration and contributes to the anagen-prolonging effects of TH seen *ex vivo*. The increased number of CD31+ endothelial cells seen, namely after low dose of topical T3 and T4 in vehicle A and T3 1nM and T4 10μM in vehicle B, further encourages this line of follow-up research, not the least since both VEGF-A and HGF are key angiogenesis promoters in human skin (Keren et al. 2022; Leung et al. 1989; Shibuya 2011; Tomita et al. 2003; Xin et al. 2001) while blood vessel regression is observed in the dermal papilla of HFs located in the balding skin of androgenetic alopecia patients (Deng et al. 2022).

If one compares all examined test groups and read-out parameters (see **Table 3**), it is challenging to identify the optimal TH dose and vehicle most likely to promote hair growth after topical administration i*n vivo*. However, acknowledging the limitations of our available preliminary pilot data, one is tempted to favor the topical application of 1 nM T3 in vehicle A since this significantly prolonged anagen *<u>and</u>* stimulated both FGF-7 production and CD31+ cell number. The latter observation raises the possibility that topical THs may even promote angiogenesis, thereby indirectly supporting anagen and thus hair growth (Mecklenburg et al. 2000; Yano et al. 2001). Yet, this remains to be formally documented by showing that topical THs indeed increase the number of CD31+ blood vessel cross sections with a lumen (Mecklenburg et al. 2000). Topically administering T3 rather than T4 also circumvents unpredictable interindividual differences that may exist in the efficacy of the intracutaneous enzymatic conversion of T4 into T3 (van Beek et al. 2008; Paus et al. 2020). Though it has been reported that DIO2, a key enzyme for the conversion of T4 to T3 is only transcribed by isolated, cultured dermal papilla fibroblasts from balding scalp (Chew et al. 2016), these cell culture results conflict with our prior finding that frontotemporal scalp skin HFs from female patients undergoing facelift surgery do express DIO2 and DIO3 (van Beek et al. 2008). Thus, it remains to be clarified definitively whether or not intact male scalp HFs from AGA-affected individuals are indeed capable of metabolizing T4 more effectively to T3 than those of individuals who are unaffected by AGA.

**Table 3.** Summary of topical T3 and T4 treatment outcomes in organ-cultured human scalp skin.

|  | Vehicle A |  |  |  |  | Vehicle B |  |  |  |  |
| --- | --- | --- | --- | --- | --- | --- | --- | --- | --- | --- |
|  | T3 1nM | T3 10nM | T4 1uM | T4 10uM | Combo | T3 1nM | T3 10nM | T4 1uM | T4 10uM | Combo |
| Hair shaft production | ↑ | ↑ | ↔ | ↑ | ↔ | ↔ | ↓ | ↓ | ↑ | ↑ |
| Anagen prolongation | ↑* | ↑ | ↔ | ↑ | ↓ | ↔ | (↑) | ↔ | ↑ | ↔ |
| HM proliferation | ↓ | ↔ | ↔ | ↓ | ↓* | ↔ | ↔ | ↓ | ↓ | ↓ |
| IGF1 | ↔ | ↓ | ↔ | ↔ | ↑ | ↑ | ↑ | ↑ | ↔ | ↑* |
| TGFB2 | ↔ | ↓ | ↔ | ↓ | ↔ | ↔ | ↔ | ↑** | ↔ | ↔ |
| FGF7 | ↑*** | ↑* | ↑*** | ↑* | ↑** | ↑ | ↑ | ↑ | ↑ | ↔ |
| Melanin content | ↔ | ↔ | ↔ | ↔ | ↔ | ↔ | ↔ | ↔ | ↔ | ↔ |
| K15 expression | ↔ | ↑** | ↑* | ↔ | ↔ | ↔ | ↔ | ↑* | ↔ | ↑ |
| K15 cell number | ↔ | ↔ | ↔ | ↔ | ↓ | ↔ | ↔ | ↑** | ↑* | ↔ |
| % of CD31+ cells | ↑* | ↔ | ↔ | ↑* | ↑ | ↑** | ↔ | ↑* | ↑ | ↑ |

Given the above considerations, it arguably may be most promising to further develop vehicle A and 1 nM T3 for subsequent clinical testing of their efficacy in managing telogen effluvium, e.g., associated with AGA. This would also circumvent the issue of interindividual and/or skin region-dependent differences in enzymatically converting T4 and T3. Before clinical testing, further vehicle optimization is advisable to reduce the risk of skin irritation upon long-term topical application of TH *in vivo* and to improve TH stability in the presence of water, e.g., by adjusting the pH and adding antioxidants. For clinical applications, the addition of viscosity-enhancing hydroxypropyl cellulose, which was necessary for our assay to avoid spillover effects of topically applied TH into the culture medium, is dispensable. An optimized vehicle should again be interrogated preclinically before it is entered into a clinical trial, using the scalp skin organ culture assay and read-outs reported here, complemented by a battery of epidermal barrier biomarkers.

In contrast to what we had seen after ‘systemic’ stimulation of isolated human scalp HFs *ex vivo* with TH or eprotirome (van Beek et al. 2008; Oláh et al. 2016), we did not observe increased HF melanin production by topically applied TH in scalp skin organ culture. Since T4 reduces the intrafollicular activity of the peripheral clock (Hardman et al. 2015a), which in turn stimulates human HF pigmentation *ex vivo* (Hardman et al. 2015b), we had expected to see a notable increase in HF melanogenesis. The duration of treatment may not have sufficed to elicit this effect, and the amount of bioactive TH that reached the HF pigmentary unit within six days may have been insufficient. It certainly would be interesting to search for subtle signs of activation of the HF pigmentary unit by topical TH that could have been missed by melanin histochemistry, for example, by assessing gp100 and MITF protein expression as well as tyrosinase activity *in situ* by IF microscopy (Gáspár et al. 2011; Hardman et al. 2015b; Suzuki et al. 2023).

Every novel pharmacological intervention for a non-life-threatening medical problem such as telogen effluvium is challenged to cautiously balance the desired clinical benefits against the risks of potential adverse effects. Given TH’s substantial but extremely well-known toxicity (Paus et al. 2020), their topical application for hair loss management purposes is no exception to this rule. Namely, the increasing evidence of potential carcinogenic effects of very long-term *systemic* T4 medication (Ho et al. 2023; Kim et al. 2024; Wändell et al. 2020) cautions against continuous, long-lasting TH administration, even if TH are applied only topically, and encourages one to test whether a pulse treatment regimen that is interspersed with long treatment-free intervals suffices to reduce telogen effluvium satisfactorily *in vivo*. Also, even though our preliminary K15 protein expression data do not support these concerns, the adverse effects of long-term topical TH administration on the bulge stem cell niche (Tiede et al. 2010) must be rigorously characterized *in vivo*.

With these caveats firmly in mind, the current proof-of-principle study introduces into the future management of hair loss disorders an innovative hormonal strategy for the treatment of hair loss disorders associated with premature anagen termination: our pilot data support the concept that telogen effluvium can be reduced by anagen-prolonging THs that can be topically applied to human scalp skin by relatively simple vehicles. This novel approach transcends, for example, the traditional inhibition of androgen receptors and dihydrotestosterone synthesis in the treatment of AGA and refocuses interest in hair loss management from steroid hormones like androgens and estrogens onto THs (Paus et al. 2020).

The 2 vehicles tested in the present study consist in a mixture of ethanol/propylene glycol/water mixture with a reduce amount of ethanol in vehicle A and greater amount of propylene glycol in vehicle B. The increase of propylene glycol a known penetration-enhancing agent should increase the flux and quantity of THs compared to vehicle A. ↔: no effect of TH treatment; ↑: stimulatory effect of TH treatment; ↓: inhibitory effect of TH treatment. *p<0.05, **p<0.01, ***p<0.001. Red rectangle indicates the optimal TH dose and vehicle most likely to promote hair growth after topical administration.

## Acknowledgements

The study was co-funded by HairDAO and CUTANEON. The authors thank Andrew Bakst for critical comments, the HairDAO community (www.hairdao.xyz) for feedback on study design in the preparatory phase and for supporting this preclinical research project. All authors are grateful to the patients who kindly donated scalp skin for this study.

## Author contributions

The study was conceived & designed by RP, and co-supervised by JC and RP, who also wrote the first manuscript draft. JG, AA, MG, AV, SV, and JC jointly generated the experimental data and interpreted the results, together with RP. UK designed the vehicles used here, and RK contributed scalp skin samples. All authors have read and edited the manuscript.

## SUPPLEMENTARY FIGURES

**Supplementary Figure 1:**
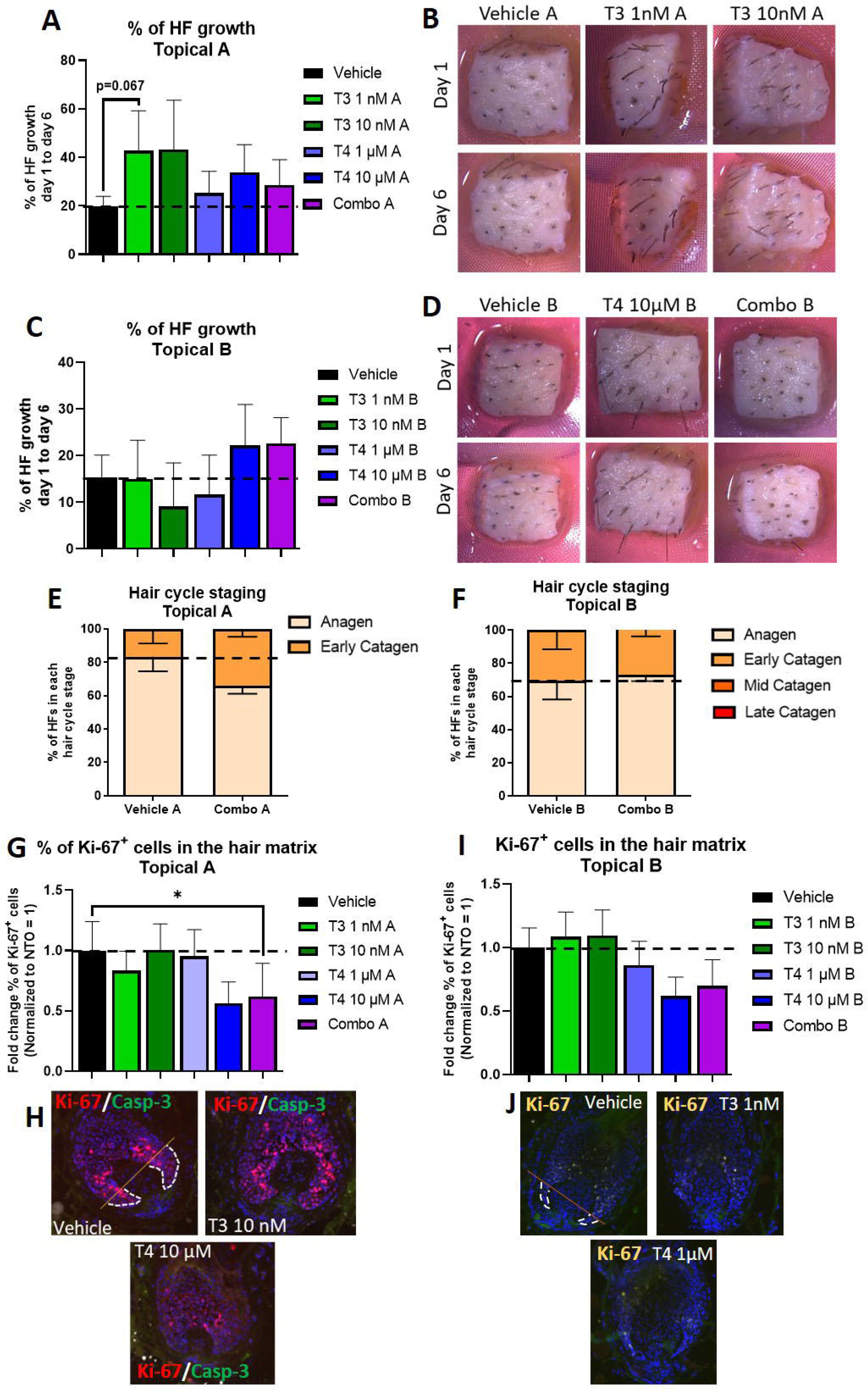
Topical thyroid hormones increase hair shaft production while the combination of T3 and T4 induces catagen and reduces hair matrix keratinocyte proliferation in human scalp skin ex vivo. **(A, C)** quantitative analysis of hair shaft production as previously described (Lu et al. 2007). N = 8-24 HFs from 1-3 different donors treated with T3 (1 or 10nM), T4 (1 or 10 µM), or vehicle. **(B, D)** Representative bright field macroscopic images of human skin fragments used for accurate hair shaft production. Mean ± SEM, Mann–Whitney test, not significant. **(E-F)** Hair cycle staging was performed using Ki-67 and Masson-Fontana histochemistry as described (Kloepper et al. 2010; Oh et al. 2016). Mean +/- SEM; n=20-35 HFs from 3 donors treated with Combo [T3 (10nM) + T4 (10 µM)], or vehicle; Mann–Whitney test, not significant. **(E, G)** qIHM of Ki-67+ cell number. Mean +/- SEM; n=17-28 HFs from 2 donors treated with T3 (1 or 10nM), T4 (1 or 10 µM), or vehicle; Mann–Whitney test, *p<0.05. (F, H) Representative images of Ki-67/Caspase-3 immunofluorescence. White dotted area: reference areas used for qIHM in the hair matrix of non-consecutive sections. Orange line: Auber’s line. Magnification: 200x.

**Supplementary Figure 2:**
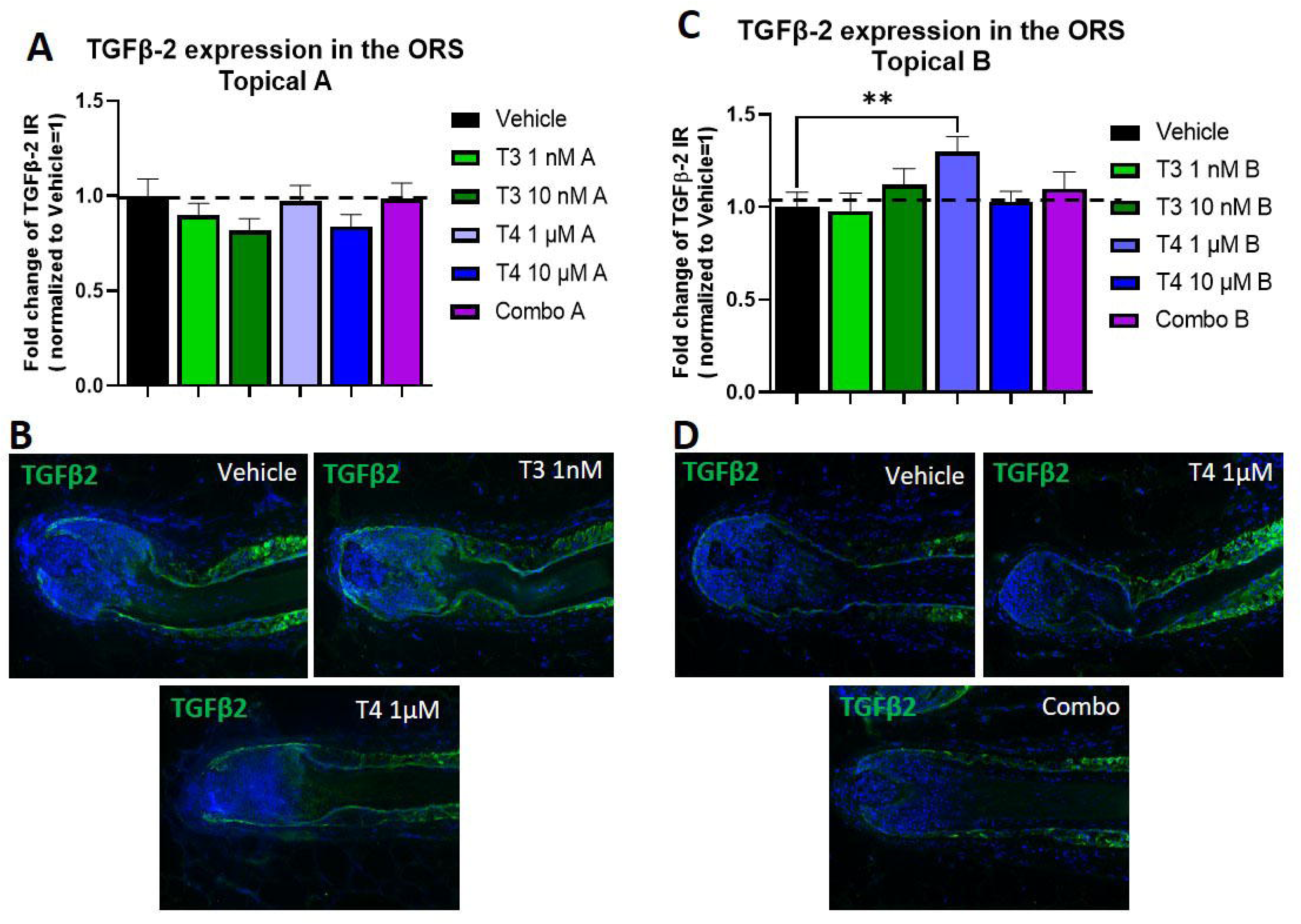
Topical thyroxine in vehicle B significantly increases TGF-β2 expression in human scalp skin *ex vivo*. **(A, C)** qIHM of TGF-β2 expression. Mean +/- SEM; n=17-33 HFs from 3 donors treated with T3 (1 or 10nM), T4 (1 or 10 µM), or vehicle; Mann–Whitney test, **p<0.01. **(B, D)** Representative images of TGF-β2 immunofluorescence. White dotted area: reference areas used for qIHM in the proximal outer root sheath of non-consecutive sections. Magnification: 200x.

